# Expression of mRNA encoding two gain-of-function *cyfip2* variants associated with DEE65 results in spontaneous seizures in *Xenopus laevis* tadpoles

**DOI:** 10.1101/2022.12.07.519540

**Authors:** Sandesh Panthi, Paul Szyszka, Caroline W. Beck

## Abstract

Developmental and epileptic encephalopathies (DEE) are a genetically diverse group of disorders with similar early clinical presentations. DEE65 is caused by *de novo*, non-synonymous, gain-of-function mutations in CYFIP2. It presents in early infancy as hypotonia, epileptic spasms and global developmental delay. While modelling loss-of-function mutations can be done using knockdown or knockout techniques to reduce the amount of functional protein, modelling gain-of-function mutations requires different approaches. Here, we show that transient ectopic expression of the Arg87Cys pathogenic variant of *cyfip2* mRNA in *Xenopus laevis* tadpoles resulted in increased seizure-related behaviours such as rapid darting and swimming in circles. In contrast, expression of a second pathological variant, Tyr108Cys, did not alter tadpole behaviour. Expression of either pathogenic variant resulted in spontaneous epileptic activity in the brain. For both variants, neuronal hyperactivity was reduced by treating the tadpole with 5 mM of the anti-seizure drug valproate (VPA). mRNA overexpression of gain-of-function variants in *X. laevis* tadpoles may be useful both for understanding the aetiology of DEE and for pre-clinical drug testing.

## INTRODUCTION

Developmental and epileptic encephalopathy (DEE) is characterised by frequent epileptiform activity that results in difficult to control seizures accompanied by developmental arrest or regression in infants and young children (Scheffer *et al*. 2017). DEE incorporates previously described syndromes of infantile onset epilepsy such as Otahara, Dravet and West syndromes, as well as later onset syndromes such as Lennox Gastaut (reviewed in (Scheffer and Liao 2020)). Children have both seizures and abnormal electrical brain activity between seizures that together result in cognitive and behavioural deterioration beyond what might be expected for the underlying pathology (Scheffer *et al*. 2017). The onset of frequent, often prolonged, uncontrollable seizures occurs on the background of normal or delayed development. Mortality is 25% by 20 years of age and survivors often have significant or profound lifelong intellectual and motor disabilities and in some cases additional sensory deficits and /or psychiatric or behavioural diagnoses (Camfield and Camfield 2008; Keezer *et al*. 2016).

Due to the increase in genome and whole exome sequencing of patients with early onset epilepsy (Epi4K *et al*. 2013), there are now more than 100 genes with nonsynonymous mutations that cause DEE reported in the Online Mendelian Inheritance in Man database (OMIM). One of the best characterised is *SCN1A*, which encodes the α2 subunit of the neuronal sodium ion channel Nav1.1. SCN1A loss of function variants cause Dravet syndrome (DEE6A, (Claes *et al*. 2001; Vadlamudi *et al*. 2010)) or the more severe non-Dravet DEE6B, which has an earlier onset and additional hyperkinetic movement disorder (Sadleir *et al*. 2017). Similar mutational spectra occur in other genes linked to DEE, which complicates diagnosis as well as the future design of therapies. Simple pre-clinical models of these de novo mutations could assist with affirming genetic diagnosis as well as provide opportunities to understand the underlying biochemistry that results in these catastrophic seizure phenotypes.

Aquatic models of epilepsy, such as *Xenopus laevis* (African clawed frog), *Xenopus tropicalis* (Western clawed frog) and *Danio rerio* (zebrafish) have already made great contributions to the understanding of specific DEE (Grone and Baraban 2015). The fish model has the advantage over the mouse model in that lines can be maintained as heterozygotes due to the tetraploid nature of *D. rerio* (Schoonheim *et al*. 2010; Cheah *et al*. 2012). In particular, study of Dravet syndrome in zebrafish *scn1ab* homozygous mutants, which exhibit seizures from 3 days post fertilisation, has led to the discovery of a novel drug target, clemizole (Baraban *et al*. 2013). Clemizole is an antihistamine, but also binds to serotonin receptors which are well conserved between fish and humans. This study has led to the identification of serotonin modulation as a target therapy for Dravet, and the initiation of human trials (Griffin *et al*. 2016). There are also zebrafish models of DEE44, caused by loss of function mutation in *UBA5*, and DEE94, caused by loss of function or haploinsufficiency at *CHD2* (Suls *et al*. 2013; Colin *et al*. 2016). CRISPR/Cas9 based knockdown in *Xenopus tropicalis* tadpoles was used to verify loss-of-function missense mutations that affect DNA binding of the transcription factor NEUROD2 as causative for DEE72 (Sega *et al*. 2019). Morpholino based knockdown and injection of DNA plasmids harbouring pathogenic missense variants of GABBR2 were used to show that reduced function of this GABA receptor in the CNS of tadpoles mimics DEE59 (Yoo *et al*. 2017). In both these studies, phenotyping was based on observation of altered tadpole swimming patterns, showing that this is a sensitive detection tool. Loss-of-function variants can therefore be modelled with knockdown/knockout strategies, with specific variants being added in as mRNA to determine if they can rescue the knockdown (Yoo *et al*. 2017, Sega, 2019 #330). However, a reliable method to verify potential gain-of-function missense pathogenic variants has not been reported before.

Cytoplasmic fragile X mental retardation protein (CYFIP) forms part of the Wiscott-Aldrich syndrome protein family verprolin-homologous protein (WAVE)-related complex (WRC), and this interaction controls neuronal connectivity via regulation of actin dynamics (Schenck *et al*. 2003). *De novo* missense mutations in *CYFIP2*, affecting Arg87, were first shown to cause DEE in four unrelated children in 2018 (Nakashima *et al*. 2018). The Arg87 site appears to be a hotspot for mutation, with 11 further patients described since(Peng *et al*. 2018; Zhong *et al*. 2019; Zweier *et al*. 2019; Begemann *et al*. 2021). Arg87 and a second mutational hotspot residue, Asp724, are located at the interface with another WRC protein, WASF1 (also called WAVE1) (Nakashima *et al*. 2018; Begemann *et al*. 2021). These CYFIP2 mutations are predicted to destabilise binding, releasing the active VCA domain of WASF1 and leading to actin polymerisation in the absence of a GDP-rac1/GTP-rac1 stimulus, resulting in the gain of function (Zweier *et al*. 2019). In patient-derived fibroblast cells, fewer dorsal ruffles (transient F actin containing ring structures that form in response to growth factors (Suetsugu *et al*. 2003)) were formed compared to controls (Begemann *et al*. 2021). Arg87 mutations are also predicted make the CYFIP2 protein less stable due to loss of electrostatic interactions with Glutamines 649 and 714 (Lee *et al*. 2019). The nearby Tyr108Cys variant, described from a single patient, is predicted to result in the loss of a phosphorylation site (Zweier *et al*. 2019). Fibroblasts derived from this patient were additionally unable to migrate properly in a wound scratch assay (Zweier *et al*. 2019).

Until very recently, no animal models of gain-of-function mutations leading to DEE65 had been reported, although two loss of function alleles (*nevermind* and *triggerhappy)* were identified from behavioural mutant screens in zebrafish (Trowe *et al*. 1996; Marsden *et al*. 2018). To accurately model the predicted gain-of-function of human DEE65 pathogenic variants that have profound neurodevelopmental disorders and intractable epilepsy, different genetic approaches are needed. In mice, Kang et al (2022) reported characteristic spasms, microcephaly, and impaired social communication in a mouse knock-in of the Arg87Cys hotspot variant *cyfip2*^+/R87C^ (Kang *et al*. 2022). Here, we show that transient expression of *cyfip2* variant-encoding mRNA in *X. laevis* tadpoles is able to both recapitulate the seizure phenotype of DEE65 and also increase agitated behaviours. Using our model, we were able to record seizure activity from the tadpole brain, and show how this is altered by the presence of the anti-seizure drug VPA.

## 2.0 MATERIALS AND METHODS

### 2.1 Preparation of mRNA for *cyfip2*-WT, Arg87Cys and Tyr108Cys pathogenic variants

Plasmid constructs of wildtype and pathogenic variant *X. laevis cyfip2.S* in pcDNA3.1 vector were obtained from Genscript. The inserts were isolated by restriction digestion with BstBI and HincII (NEB) and sub-cloned into CS2^+^ vector cut with BstB1 and StuI (NEB) using T4 ligase (Roche). Plasmids were verified by sequencing and purified, before cutting with NotI to linearise. The resulting linear DNA templates were *in vitro* transcribed into capped mRNA using a mMessage mMachine Sp6 kit (Ambion). mRNA was stored at −80°C in 2 μl aliquots. Capped *nGFP* mRNA was prepared from CS2^+^ nucGFP in the same way.

### 2.2 Husbandry and production of *X. laevis* eggs and embryos

*Xenopus laevis* male and female frogs were bred at the University of Otago’s Zoology department and housed in a recirculating Marine Biotech XR3 aquarium maintained at 18°C with a 12/12 hour light/dark cycle. Adult female *X. laevis* were induced to lay eggs by injecting human chorionic gonadotrophin (Chorulon) (500 IU per 75 g body weight) into the dorsal lymph sac. Injections were performed ~16 hours before eggs were required. Next morning, each female was placed in 1 L of 1x Marc’s modified ringer’s solution (MMR: 100 mM NaCl, 2 mM KCl, 1 mM MgSO_4_, 2 mM CaCl_2_, 5 mM HEPES, 0.1 mM EDTA pH 8.0) Eggs were then collected hourly and fertilised using fresh testes from a euthanised adult *X. laevis* male. Fertilised eggs were treated with 2% w/v cysteine solution (pH 7.8) to remove the jelly coats, and rinsed three times in 1x MMR. Embryos were staged according to Nieuwkoop and Faber’s staging series (Nieuwkoop and Faber 1994).

### 2.3 Micro-injection of capped mRNA into *X. laevis* embryos

Fertilised eggs were identified by the presence of a sperm entry point and placed in wells cut into 2% noble agar lined 50 mm petri dishes containing 5% Ficoll 4000 (Sigma) in 1xMMR. Each embryo was injected with 9.2 nl of mRNA (500 pg of nGFP and 4 ng of one of the *cyfip2* mRNAs) using a backfilled capillary needle and Drummond Nanoject II microinjector. Media was replaced with 3% Ficoll in 0.1 x MMR at mid blastula stage before overnight incubation at 18°C. Healthy embryos were transferred to 0.1x MMR in 10 cm petri dishes and raised to stage 46.

### 2.4 Behavioural phenotyping of tadpoles

Individual tadpoles at stage 47 were arrayed in a 24-well plate (Greiner Bio-one cellstar) containing 1.2 ml of 0.1 x MMR. The movement of tadpoles was then recorded from above using a Panasonic DMC-FZ1000 set in Movie mode (25 frames per second, MP4), with strong diffuse LED lighting from below. Tadpole behaviour was recorded for 90 minutes. The video was converted into MPEG2 format using Adobe Media Encoder CC 2019 software. Mean swimming velocity and percentage of time spent circling or darting was automatically calculated using TopScan (CleverSys) as described in (Panthi *et al*. 2022).

### 2.5 Simultaneous local field potential (LFP) recording from optical tectum and muscle

The most active tadpoles from video analysis were selected for electrophysiology to detect seizure activity. Tadpoles at stage 47 were first briefly immersed in 0.5 mM of the neuromuscular blocker pancuronium bromide (Sigma) on a Sylgard-lined petri dish. Once movement had ceased, 4-5 minutien needles were placed around the head and trunk to stabilise the tadpole. A small incision was made in the dorsal skin of the head to expose the underlying optic tectal lobes, using a piece of razor blade as a knife. Tadpoles were transferred to a 2.5 cm petri dish and surrounded with cooled 2% low melting point agarose to adhere them to the plate with the brain facing upwards. The dish was then flooded with HEPES-buffered extracellular saline (115 mM NaCl, 4 mM KCl, 3 mM CaCl_2_, 3 mM MgCl_2_, 5 mM HEPES, 10 mM glucose, pH 7.25). The recording electrodes were HEPES-buffered extracellular saline-filled glass micropipettes (type 1B100F-3, World Precision Instruments) pulled with a Sutter P-87 micropipette puller, to get a tip resistance of approximately 10 mΩ. Two electrodes were mounted in micromanipulators (Kerr Scientific Instruments Tissue Recording System) and positioned so that one was in the optic tectum (brain signals) and the other in the tail muscle (muscle signals). Brain and muscle signals were differentially amplified against a common reference electrode (silver wire of 0.27mm diameter) that was submerged in the saline bath of the tadpole recording chamber using 250x gain, AC-coupling and 2 kHz low pass filter (MA 103 preamplifier and 102 4-channel amplifier, Universität zu Köln). Noise from the powerline (50 Hz) was reduced with a Hum Bug (Quest Scientific), and signals were digitised at 10 kHz (Micro 3 140, Cambridge Electronic Design). All recordings were performed at room temperature (20–22°C), under daylight, on an anti-vibration table and inside a Faraday cage.

Power spectrum analysis and spike sorting was carried out using Spike2 v8 (Cambridge Electronic Design). Power spectra were calculated with Hanning windows and an FFT size of 262144. For each tadpole, power spectra were calculated for 25 minutes long signal traces. Before spike sorting, the optic tectum signal was digitally high-pass filtered (>0.1 Hz) and down-sampled to 1 kHz. Spikes were first detected automatically, and then reviewed manually. A spike was defined as a negative or positive deflection greater than 5 or 10 times the standard deviation of baseline activity (a 15 minutes long period of activity with no visible spikes). The “spike window” was adjusted for each tadpole depending on how rapid the spikes were, from 50 ms. A spike that was mirrored in the muscle signal was excluded if the spike in the muscle signal was > 0.2 x the amplitude of the corresponding spike in the optic tectum. As positive controls, recordings were also made from tadpoles treated with 5 mM pentylenetetrazole (PTZ), to induce acute seizures. Representative traces were exported from Spike2 as .emf graphics.

### 2.6 Statistical analysis

All data were graphed and analysed using Prism v9.0. Raw data and statistical analysis results can be found in the supplemental data file.

## 3.0 RESULTS

### 3.1 *X. laevis cyfip2* is expressed in tadpole brains and the protein is almost identical to human CYFIP2

*X. laevis* is an allotetraploid and there are both S and L homeologues of *cyfip2* in the genome, which code identical proteins. *Cyfip2.S* is expressed at much higher levels during development (Session *et al*. 2016) and is high in the tadpole midbrain from stage 44 to 61 according to data from Ta et al. (Ta *et al*. 2022), Fig S1). We therefore used the *cyfip2.S* sequence to recreate two human variants associated with more severe forms of DEE65, Arg87Cys and Tyr108Cys. The *X. laevis* protein is 98% identical to human CYFIP2, and the amino acids at sites of interest are conserved (Fig. 1A, Fig. S2). Alphafold rendering of the *X. laevis* Cyfip2 protein is almost indistinguishable from human, with the positions of the two pathogenic variants located in the same pocket, identified as binding WAVE1 VCA domain by Zweier et al (Zweier *et al*. 2019).

**Figure 1:**
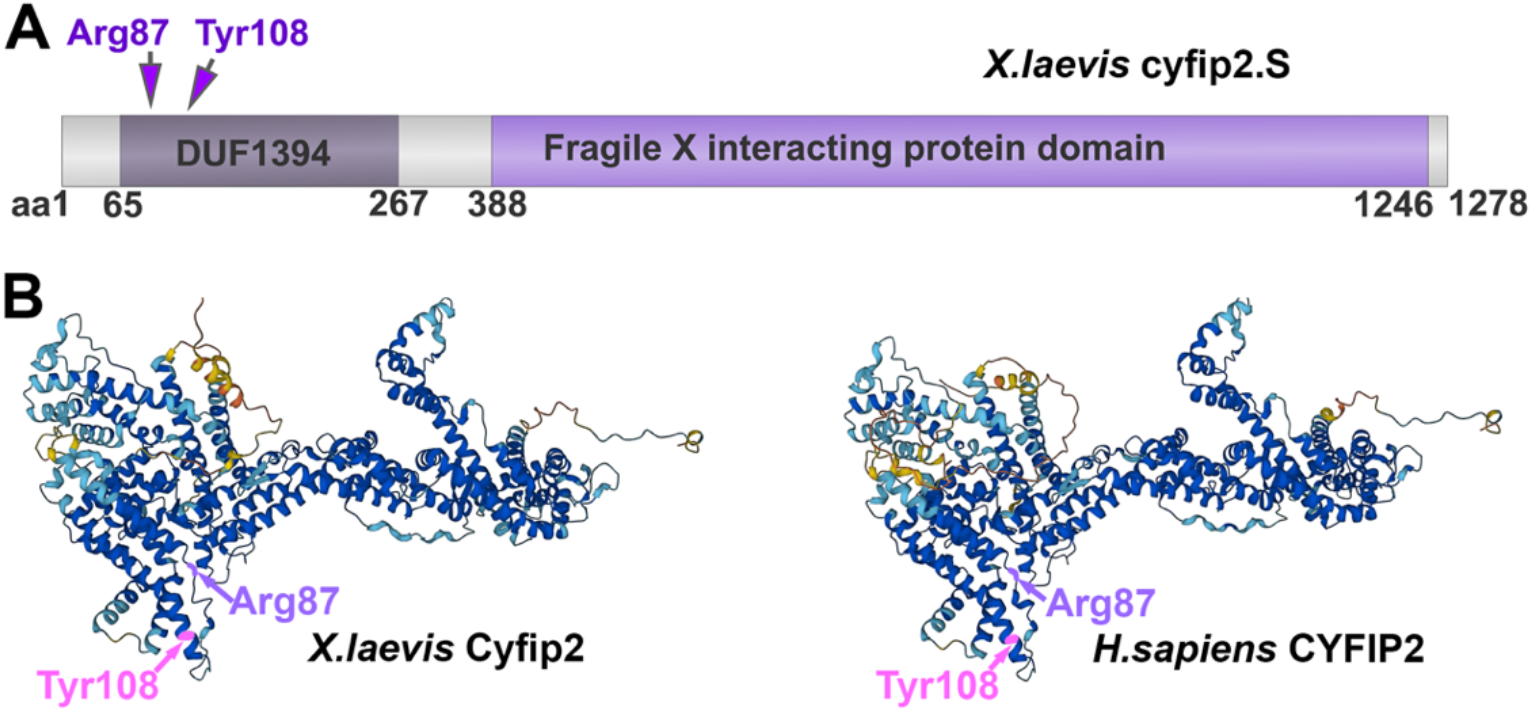
*X. laevis* Cyfip2 is conserved at variant sites and predicted protein structure is very similar to human. A) Schematic of the protein domains of Cyfip2 from *X. laevis* showing the locations of the two most severe DEE65 causing variants. DUF is domain of unknown function, but Arg87 and Tyr108 are located in a pocket that binds the VCA domain of WAVE1. B) Alphafold protein structure predictions of *X. laevis* (left) and human (right) proteins showing the locations of the variants (Jumper *et al*. 2021).

### 3.2 Embryonic injection of mRNA encoding the pathogenic variant *cyfip2-*R87C induces seizure like behaviours

Tadpoles overexpressing *cyfip2-*WT or the pathogenic variants Arg87Cys and Tyr108Cys appeared morphologically normal at stage 47 (pre-feeding stages). Behavioural analysis of five tadpole sibships was extracted by TopScan from 90 minutes of video recordings of individual tadpoles arrayed in 24 well plates (Fig. 2). *Cyfip2*-R87C tadpoles were consistently and significantly more active across all three measurements, swimming velocity, percentage of time spent swimming in circles, and percentage of time spent darting. This cohort swam on average 1.4 fold faster than uninjected controls. The percentage of time spent in circling behaviours was 1.7 times and time spent darting was 1.5 times that of uninjected controls. In contrast, the tadpoles expressing *cyfip2*-Tyr108Cys did not show any obvious behavioural changes. Unexpectedly, *cyfip2* WT overexpression showed a significant increase in time spent darting. To control for any effect of co-injected nGFP mRNA, we also included tadpoles only injected with nGFP, and these behaved no differently to uninjected controls (Fig. 2). C-shaped contractions as described by Sega et al for *NeuroD2* DEE72 tadpoles models and for tadpoles chemically induced to seizure as described by Hewapathirane et al were not observed (Hewapathirane *et al*. 2008; Sega *et al*. 2019).

**Figure 2.**
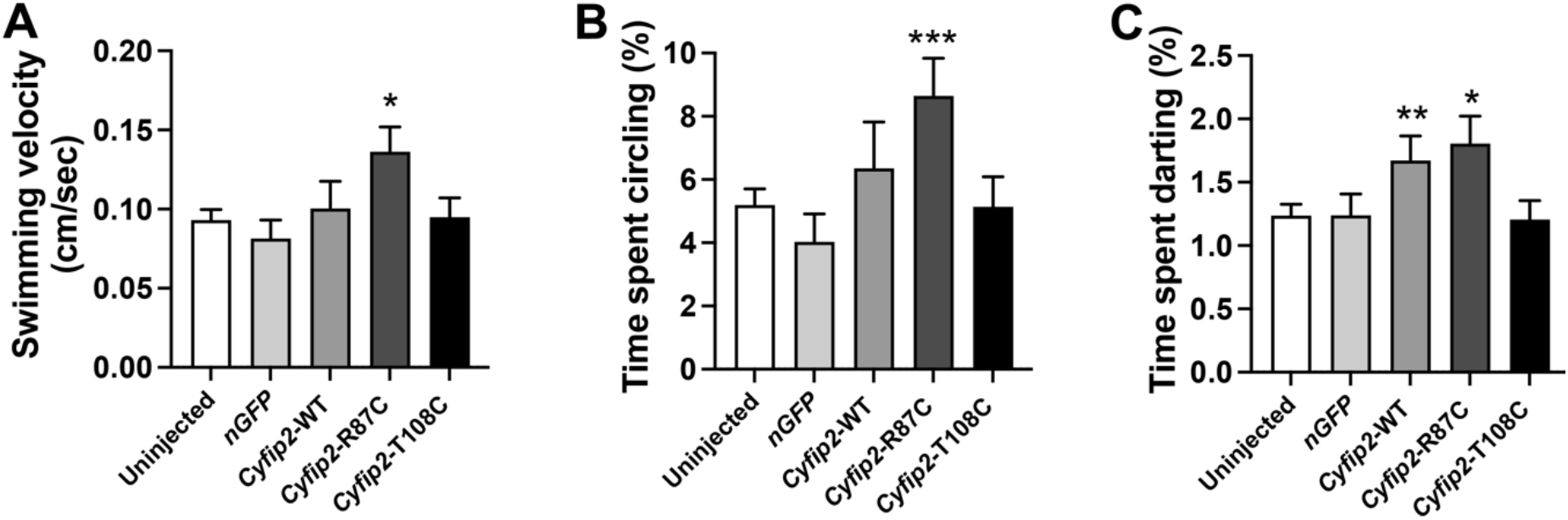
Behavioural analysis of tadpoles injected with mRNA encoding pathogenic variants of Cyfip2. A-C) Bar graphs showing mean, error bars are standard error. Tadpoles overexpressing c*yfip2*-R87C variant swim faster and spend more time in circling and darting behaviours. *Cyfip2-*WT tadpoles also spent more time darting than controls. Uninjected N=395, *nGFP* mRNA injected N=72, *cyfip2*-WT mRNA injected N=72, *cyfip2*-R87C injected N=108, *cyfip2* T108C injected N=137. Kruskal-Wallis test with Dunn’s multiple testing to control uninjected: * indicates p<0.05, ** p<0.01. Raw data can be found in the supplemental data file.

### 3.3 Local field potential (LFP) recording in the brain confirms neuronal seizure induction by PTZ

The most common test to confirm a suspected epilepsy diagnosis in humans is to record brain activity. Therefore, to confirm that the tadpoles’ seizure-like behaviours reflect epileptic seizures, we recorded neuronal activity in the brain (optic tectum) with LFP recordings. In order to ensure that electrophysiological recordings from the optic tectum were consistent with seizure activity, we first optimised recording in wild type tadpoles treated with the epileptogenic chemical pentylenetetrazol (PTZ) (Fig.3). To separate brain activity from muscle activity, we recorded from both the tail muscle and optic tectum of the same animal using two electrodes. Tadpoles were briefly immersed in the neuromuscular blocker pancuronium bromide and mounted in 2% low melting point agarose to prevent movements that would dislodge the electrodes. Due to the small size of the tadpole, some electrical activity of the muscle is also picked up in the brain and vice versa. To exclude muscle activity, we only analysed recordings where brain LFP signals were at least five times larger than muscle LFP signals.

**Figure 3.**
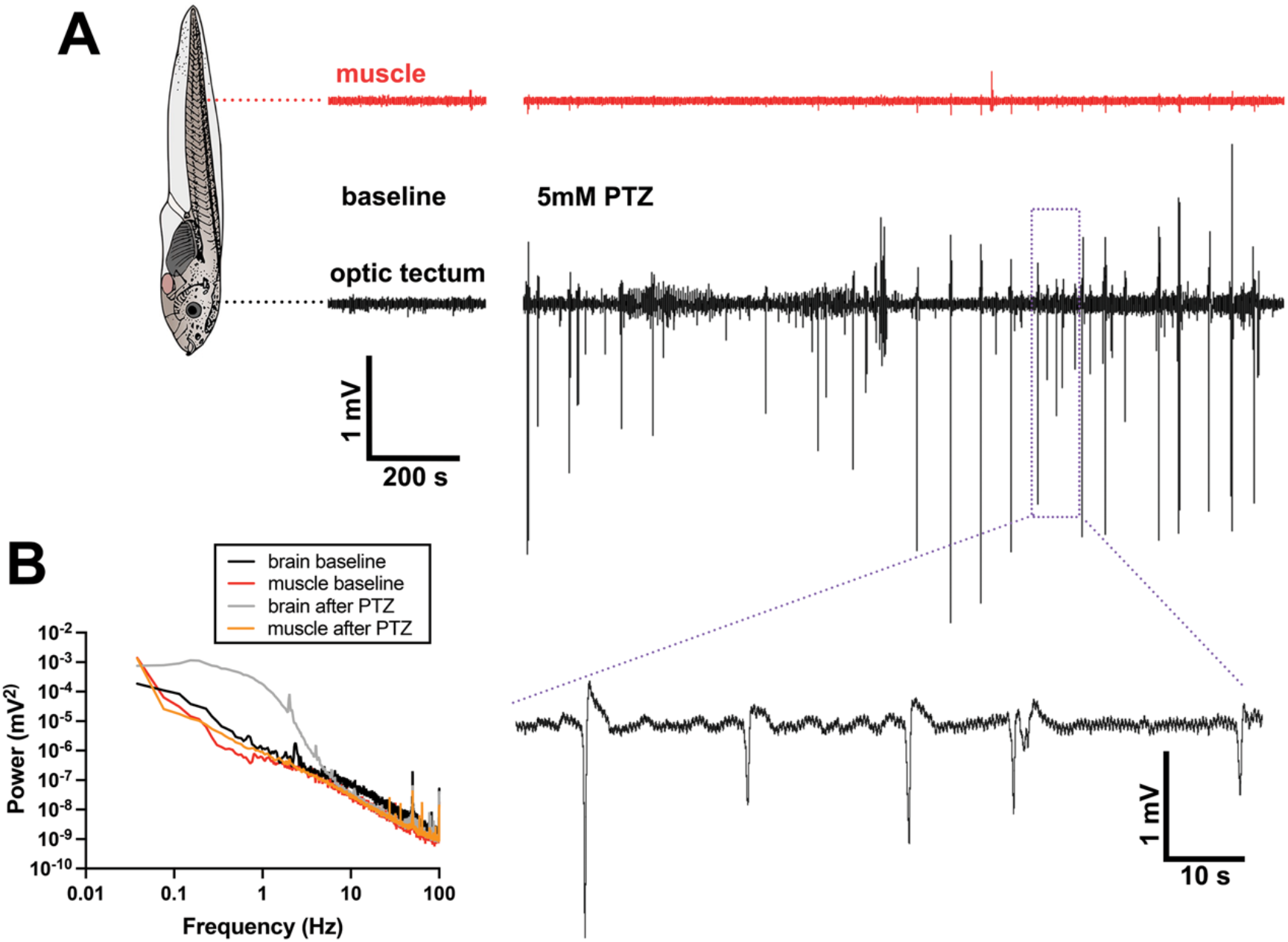
Local field potential recordings in the optic tectum can be used to detect neuronal seizures. A) LFP recordings from the optic tectum and muscle of the same tadpole, shown at the same scale. The baseline shows five minutes of recording from both channels before 5 mM PTZ application, and the trace after the break shows 30 minutes of acute PTZ-induced seizure activity. Purple dotted rectangle shows the area expanded below to show individual LFP spikes seen in the optic tectum channel. B) Power spectrum analysis of the same animal. The largest power increase in the brain following PTZ addition is in the 0.1-10 Hz range.

Adding 5 mM PTZ to the tadpole’s medium resulted in spikes of around five seconds duration and an amplitude of 4.8 mV or less (Fig. 3A) and an increase in power between 0.1 and 10 Hz in the optic tectum but not muscle (Fig. 3B). Therefore, PTZ increases the neuronal synchrony that is characteristic of seizures as shown in a previous study using the same age *X. laevis* tadpoles (Hewapathirane *et al*. 2008).

### 3.4 Spontaneous neuronal seizures can be detected in the brains of tadpoles that express pathogenic *cyfip2* variants

To test whether the seizure-like behaviours in variant *cyfip2* expressing tadpoles reflect neuronal seizures, we made LFP recordings from muscle and brain in tadpoles that had shown high swimming activity in the behavioural tests (Fig. 4). LFP recordings were also made in tadpoles from three different control groups: (i) wild type uninjected tadpoles, (ii) tadpoles from embryos injected with *nGFP* mRNA only and (iii) tadpoles expressing the wild type *cyfip2* mRNA (Fig. 4 A,D). DEE65 results in chronic seizures and so these models would be expected to have regular seizure like episodes interspersed with periods of baseline activity. We did not detect brain-specific spikes of activity in tadpoles that had not been injected with any mRNA as embryos (negative controls), nor from *nGFP* or *cyfip2-*WT mRNA expressing tadpoles. This suggests that overexpression of *Xl*.*cyfip2* is not inherently harmful to tadpoles, despite the darting behaviour increase noted in Fig. 2, and that both the injection process itself, and the presence of nGFP, are similarly benign.

**Figure 4:**
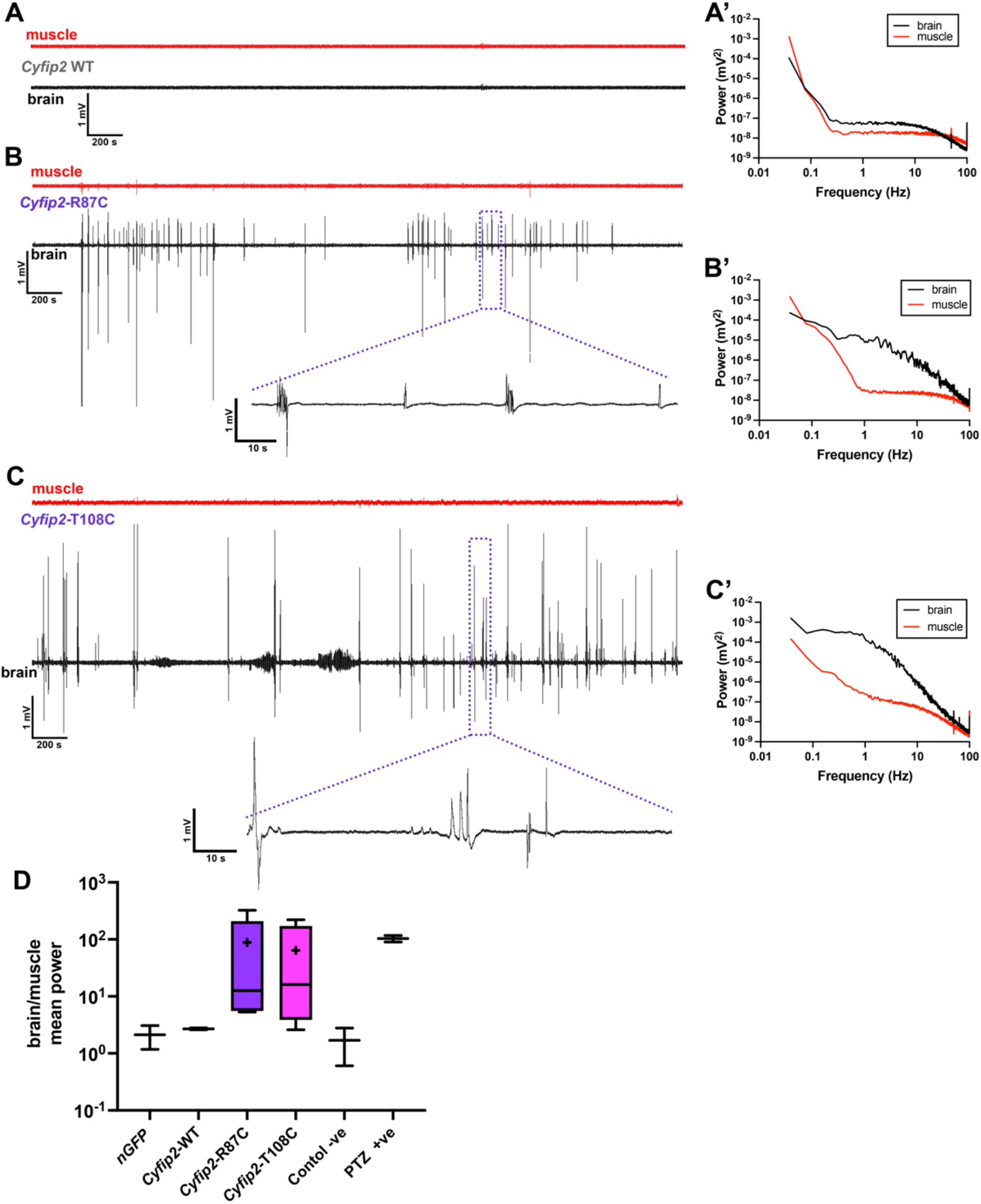
Paired brain and muscle LFP recordings reveal spontaneous neuronal seizures in tadpoles that express pathogenic *cyfip2* variants. A-C) Representative LFP signals for 60 minutes of simultaneous recording from muscle (red) and brain (optic tectum, black) of the same tadpole. Pathogenic variants are in purple text. Purple dotted boxes indicate the location of the zoomed trace portion underneath, which shows 100 seconds of brain recording for the two pathogenic variants. A) No spikes were detected in tadpoles overexpressing wild type *cyfip2*. B) LFP spikes in tadpoles expressing *cyfip2-* R87C. C) Tadpoles expressing *cyfip2-*T108C also showed a variety of distinct spikes. A’-C’ Power spectrum analysis of brain (black) and muscle (red), shown from 0-100 Hz and compiled from a 25 minute section of each trace. D) Box plot of brain:muscle ratio of mean power across 0.1-10Hz, for each group. N=2 tadpoles for *nGFP*, *cyfip-*WT, Control - ve and 5mM PTZ +ve, N= 5 for *cyfip2-*R87C, N=4 for *cyfip2-*T108C. Positive and negative controls are the same tadpoles before and after PTZ addition. + indicates mean, bar indicates median, box indicates interquartile range and whiskers indicate min/max values. All power spectra are shown in Fig. S4, and Raw data can be found in the supplemental data file.

Five tadpoles expressing the *cyfip2-*R87C variant showed brain-specific, spontaneous LFP activity. The LFP spikes were of very short duration, 1 to 1.5 seconds (Fig. 4B), and occurred as short spike clusters or single spikes with normal baseline activity between. Larger fast spikes had an amplitude up to 5 mV, similar to those elicited with 5 mM PTZ. The oscillatory power of brain activity was increased between 0.1 and 10 Hz (Fig. 4B’, D and Fig.S4), mirroring the PTZ-induced increase (Fig. 3B). Muscle activity was consistently lower over the same period (Fig. 4B, B’ and Fig.S4) showing that LFP signals from the optic tectum reflect brain-specific activity. Similarly, the *cyfip2-*T108C variant (4 tadpoles) showed spontaneous brain-specific spikes interspersed with periods of baseline activity (Fig. 4C). Spikes had different amplitudes, up to a maximum of 5 mV, as well as different shapes, and a duration of 4-6 seconds. Amplitudes also varied, but were normally <5 mV and often much smaller. Oscillatory activity was similar to that seen with PTZ or *cyfip2*-R87C (Fig. 4C’). The mean power (0.1-10 Hz) of each tadpole was calculated for both channels, and brain power divided by muscle power was graphed for all groups (Fig. 4D). Brain was an average of 88 times more active in this frequency range than muscle for *cyfip2-*R87C variants (range 5 to 324 times). For *cyfip2-*T108C tadpoles, brains were on average 63 times more active on this range than muscle (range 2.6-218 times).

To confirm the results of the power analysis, we also quantified the spike rate and inter-spike intervals (Fig. 5A,B). The highest spike rates were seen in wild type tadpoles treated with 5 mM PTZ (8.7 and 3.5 spikes/minute). *Cyfip2*-WT expressing tadpoles, in contrast, had 0.06 and 0.04 spikes/minute, which were only just above background, and *cyfip2-*R87C tadpoles averaged 0.85 ± 0.3 spikes/min and *cyfip2-*T108C 0.92 ± 0.4 spikes/min (Fig. 5A). Therefore, tadpoles that have spontaneous seizures resulting from expression of either pathogenic variant of *cyfip2* had around 17 times higher spike rates than tadpoles expressing the equivalent level of WT *cyfip2*, and a 6-7 times lower spike rate than the PTZ acute seizure model (Fig. 5A). Accordingly, inter-spike intervals were longest in the *cyfip2* WT expressing tadpoles (502.62 and 756.14 seconds). *Cyfip2-*R87C expressing tadpoles had a mean inter-spike interval of 117.29 ± 48.48 seconds and the *cyfip2-*T108C tadpoles had an average inter-spike interval of 47.33 ± 4.3 seconds (Fig. 5B). The inter-spike interval for acute seizures caused by PTZ was shortest (7.00 and 16.74 seconds).

**Figure 5:**
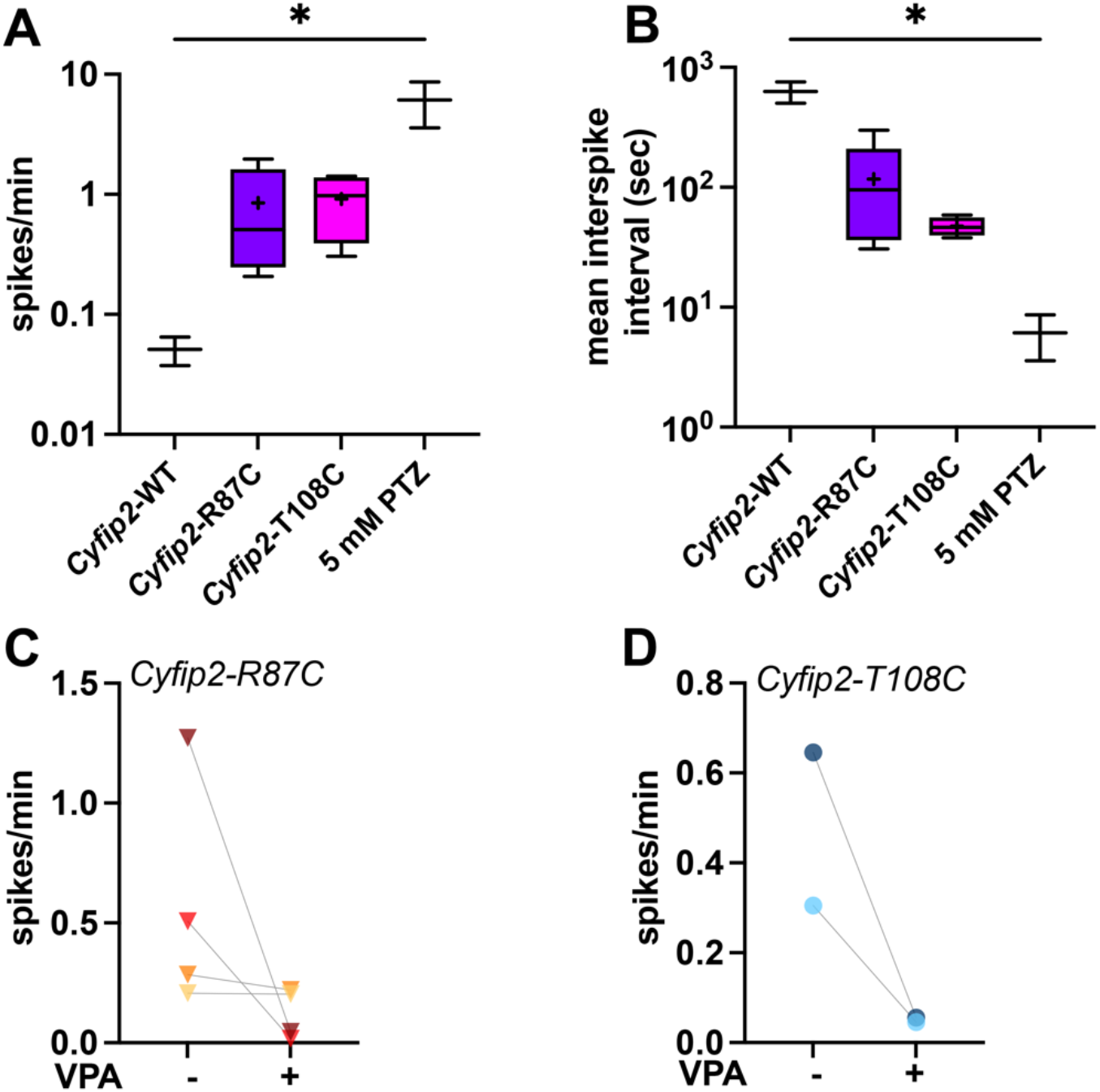
Expression of pathogenic variants of *cyfip2* increases the rate of spontaneous brain LFP spikes, and adding the antiepileptic drug VPA decreases the spike rate. **A,B** Box plots, + indicates mean, middle line indicates median, box indicates interquartile range and whiskers indicate min/max values. A) The rate of brain-specific (epileptic) spikes/min in tadpoles expressing WT *cyfip2* or one of the two pathogenic variants, compared to tadpoles treated with PTZ. B) mean inter-spike interval for WT or variant *cyfip2* compared to 5mM PTZ treated wild type tadpoles. N=2 for *cyfip2-*WT and PTZ treated tadpoles, N=5 for *cyfip2-*R87C and N=4 for *cyfip2-*T108C. Kruskal-Wallis with Dunn’s post hoc testing of all means, * indicates p<0.05. **C,D)** Before/after plots comparing spikes per minute before and after adding 5 mM VPA. Individual tadpoles expressing *cyfip2-*R87C (C) are or *cyfip2-*T108C (D) are connected by grey dotted lines, and indicated by different shading. Raw data can be found in the supplemental data file.

### 3.5 Seizure activity in *cyfip2-*R87C and *cyfip2-*T108C tadpoles is reduced in the presence of the anti-seizure drug valproic acid (VPA)

To see if our gain-of-function model for DEE65 would be useful in drug testing, we added 5mM VPA to the tadpole buffer during electrophysiological recording and compared the rate of brain-specific spikes before and after drug treatment. VPA reduced the rate of spikes in all tadpoles, although the effectiveness varied (Fig. 5 C,D). VPA reduced seizure spike rate to 14% of the pre-treatment rate for *cyfip2-*R87C variant (n=4) and to 6% of pre-treatment rates for *cyfip2-*T108C (n=2). Although the numbers of tadpoles was too small for statistical testing, these results suggest that VPA is an effective drug to reduce spontaneous seizures caused by either variant.

## DISCUSSION

In humans, DEE65 results from *de novo* mutations in *CYFIP2*, and mutations at Arg87 or Tyr108 are associated with profound neurodevelopmental and epileptic phenotypes. These variants appear to result in a gain-of-function due to the altered binding to the VCA domain of WASP1, and in fibroblasts, this leads to dysregulation of actin dynamics (Zweier *et al*. 2019). Additionally, these mutations may also lead to destabilisation of the CYFIP2 protein (Lee *et al*. 2019; Zweier *et al*. 2019).

Here, we have used mRNA overexpression of pathogenic *cyfip2-*Arg87 and *cyfip2*-Tyr108 variants during early development to determine the effect on the developing tadpole brain. We show that expression of the hotspot *cyfip2*-R87C pathogenic variant causes tadpoles to develop more agitated swimming behaviours, swimming faster than controls and spending more time in circling and darting behaviours. This behavioural hyperactivity is consistent with epileptic seizures (Hewapathirane *et al*. 2008; Panthi *et al*. 2022), and it is in line with the seizure-like behaviour (hyperactivity and epileptic spasms) recently found in a mouse knock-in model of *cyfip2^+/R87C^* (Kang *et al*. 2022). Our study is complementary to that of Kang et al (2022). In the mouse model, spontaneous seizure detection was not possible in the newborn mice, and was not seen in juveniles, but more severe seizure responses were observed when adults were exposed to pentylenetetrazole (PTZ) (Kang *et al*. 2022). In our tadpole model, we could confirm the epileptic nature of spontaneous *cyfip2-*R87C-induced seizures in electrophysiological recordings of electrical brain activity. Further, we were also able to detect spontaneous seizures in tadpoles expressing a second patient pathogenic variant, Tyr108Cys. Both of these models accurately represent the gain of function, *de novo* profound DEE65, and can be used in future for testing anti-seizure drugs and devising therapies. Additionally, we tested the effect of the anti-seizure drug VPA and showed that this reduced the rate of seizure spikes in tadpoles expressing either pathogenic variant. It is not yet known if early seizure control can change the developmental trajectory of children with DEE65, but recent reports have suggested that the GABA transaminase inhibitor vigabatrin may be a good choice to control infantile spasms (infant with Arg87Leu), (Zhong *et al*. 2019)). Cannabidiol (CBD) is now approved for use in refractory epilepsy such as Dravet and Lennox-Gastaut syndrome (Ortiz *et al*. 2022; Pagano *et al*. 2022), both of which now fall under the DEE umbrella diagnostic term. There has been one report of a three year old child with DEE65 (R87C) becoming seizure free with CBD (de Goes *et al*. 2022). CBD does not activate CB1 or CB2 receptors; instead they are thought to interact with ion channels and G-protein coupled receptors directly (Oz *et al*. 2022).

The phenotypic spectrum of DEE65 ranges from profound neurodevelopmental disability with intractable epilepsy at the hotspot Arg87, to a more variable neurodevelopmental phenotype in other characterised missense variants (Begemann *et al*. 2021). In contrast, heterozygous nonsense variants predicted to result in *CYFIP2* loss-of-function had milder symptoms and did not fit the profile of a DEE (Begemann *et al*. 2021). We have focussed here on modelling the two most severe variants of DEE65, predicted to cause aberrant actin polymerisation via altered interactions between CYFIP2 and WAVE1 (Zweier *et al*. 2019). These mutations may also destabilise CYFIP2, as suggested by (Lee *et al*. 2019) and (Kang *et al*. 2022). *Cyfip2*^−/−^ mice die perinatally (Zhang *et al*. 2018), while heterozygotes (*cyfip2*^+/−^) have increased seizure susceptibility as adults, arising from increased f-actin, enlarged dendritic spines, and enhanced excitatory synaptic transmission in prefrontal cortex neurons (Lee *et al*. 2020). Supporting this, zebrafish loss-of-function mutants of *cyfip2* have been found in behavioural mutagenesis screens. *Nevermind (nev)* was first identified from the forward genetic screens of the 1990s and classified as affecting retinotectal projections and axon sorting (Trowe *et al*. 1996). The *nev* mutation causes Cyfip2 protein truncation after just 79 amino acids, and fish die at eight days old after exhibiting abnormal swimming and failing to inflate a swim bladder. A second zebrafish *cyfip2* allele, *triggerhappy*, was identified from a screen for heightened acoustic startle response driven by Mauthner cell activation (Marsden *et al*. 2018). *Triggerhappy* results in a Cyfip2 protein truncated at amino acid 342. The startle response of fish involves a C-shaped bend which forms the first part of a “turn and run” startle response controlled by Mauthner cells which are also found in tadpoles (Sillar and Robertson 2009). Mauthner cells are coincident detectors (they generate action potentials when receiving synchronous synaptic inputs) and they are normally not spontaneously active (Korn and Faber 2005). Therefore, the seizure-related increase in neuronal synchrony seen in tadpole brains after PTZ application (Hewapathirane *et al*. 2008) and Fig. 3) could spontaneously activate Mauthner cells which would lead to C-shaped contractions seen in acute seizure models (Hewapathirane *et al*. 2008; Panthi *et al*. 2022). We did not observe C-shaped contractions in tadpoles expressing *cyfip2-*R87C, but we did show increased darting behaviour, a series of startles/rapid C-start movements leading to rapid changes in direction.

A transgenic line expressing wild type *cyfip2* under heat shock from 30 hours post fertilisation was able to rescue the abnormal startle response of *triggerhappy* fish, confirming the underlying nonsense mutation as loss-of-function (Marsden *et al*. 2018). Here, we were able to inject high levels of wild type *X. laevis cyfip2* mRNA with no adverse effects on gross phenotype, survival, or seizure activity, but we did observe increased darting behaviour. Taken together with the human data (Begemann *et al*. 2021), this suggests that while both low and high levels of Cyfip2 can affect behavioural thresholds, only predicted gain-of-function mutations result in spontaneous epileptic seizure activity. A recent study used CRISPRa system to develop a *Scn1a* zebrafish model (sodium ion channel) which showed an epileptic phenotype arising from overexpression in the absence of mutation (Weuring *et al*. 2021). Like ours, this study quantified locomotor burst movements/hyperactivity as seizure-related behaviour. In this case, the gain-of-function resulted from overexpressing the ion channel gene, which would model gene duplications, increased protein or transcript stability (Weuring *et al*. 2021). Studies in animal models of DEE can therefore confirm pathogenic mutations as seizure causing, and contribute to our understanding of genotype-phenotype correlations in this genetically complex group of neurodevelopmental disorders.

## Supporting information

raw data

## ACKNOWLEDGEMENTS

The authors would like to thank Nikita Woodhead for excellent care of the *Xenopus* colony, Joanna Ward for laboratory assistance, Sulagna Banerjee for assistance with TopScan, and Dr. Phoebe Chapman and Cabriana Earl for critical reading of the manuscript. This work was funded by the Neurological Foundation of New Zealand (Project Grant 2025PRG to CWB, PS and Prof. Lynette Sadlier).

## DATA AVAILABILITY

Plasmids are available upon request. The authors affirm that all data necessary for confirming the conclusions of the article are present within the article, figures, supplementary figures, and raw data can be found in the supplemental data file.

## REFERENCES

Baraban, S. C., M. T. Dinday and G. A. Hortopan, 2013 Drug screening in Scn1a zebrafish mutant identifies clemizole as a potential Dravet syndrome treatment. Nat Commun 4: 2410.

Begemann, A., H. Sticht, A. Begtrup, A. Vitobello, L. Faivre et al., 2021 New insights into the clinical and molecular spectrum of the novel CYFIP2-related neurodevelopmental disorder and impairment of the WRC-mediated actin dynamics. Genet Med 23: 543–554.

Camfield, C., and P. Camfield, 2008 Twenty years after childhood-onset symptomatic generalized epilepsy the social outcome is usually dependency or death: a population-based study. Dev Med Child Neurol 50: 859–863.

Cheah, C. S., F. H. Yu, R. E. Westenbroek, F. K. Kalume, J. C. Oakley et al., 2012 Specific deletion of NaV1.1 sodium channels in inhibitory interneurons causes seizures and premature death in a mouse model of Dravet syndrome. Proc Natl Acad Sci U S A 109: 14646–14651.

Claes, L., J. Del-Favero, B. Ceulemans, L. Lagae, C. Van Broeckhoven et al., 2001 De novo mutations in the sodium-channel gene SCN1A cause severe myoclonic epilepsy of infancy. Am J Hum Genet 68: 1327–1332.

Colin, E., J. Daniel, A. Ziegler, J. Wakim, A. Scrivo et al., 2016 Biallelic Variants in UBA5 Reveal that Disruption of the UFM1 Cascade Can Result in Early-Onset Encephalopathy. Am J Hum Genet 99: 695–703.

de Goes, F. V., J. T. M. de Andrade Ramos, R. da Silva Fontana, C. L. de Carvalho Serao, F. Kok et al., 2022 Cannabidiol Successful Therapy for Developmental and Epileptic Encephalopathy Related to CYFIP2. The Open Neurology Journal 16.

Epi4K, P. Epilepsy Phenome/Genome, A. S. Allen, S. F. Berkovic, P. Cossette et al., 2013 De novo mutations in epileptic encephalopathies. Nature 501: 217–221.

Griffin, A., C. Krasniak and S. C. Baraban, 2016 Advancing epilepsy treatment through personalized genetic zebrafish models. Prog Brain Res 226: 195–207.

Grone, B. P., and S. C. Baraban, 2015 Animal models in epilepsy research: legacies and new directions. Nat Neurosci 18: 339–343.

Hewapathirane, D. S., D. Dunfield, W. Yen, S. Chen and K. Haas, 2008 In vivo imaging of seizure activity in a novel developmental seizure model. Exp Neurol 211: 480–488.

Jumper, J., R. Evans, A. Pritzel, T. Green, M. Figurnov et al., 2021 Highly accurate protein structure prediction with AlphaFold. Nature 596: 583–589.

Kang, M., Y. Zhang, H. R. Kang, S. Kim, R. Ma et al., 2022 CYFIP2 p.Arg87Cys Causes Neurological Defects and Degradation of CYFIP2. Ann Neurol.

Keezer, M. R., G. S. Bell, A. Neligan, J. Novy and J. W. Sander, 2016 Cause of death and predictors of mortality in a community-based cohort of people with epilepsy. Neurology 86: 704–712.

Korn, H., and D. S. Faber, 2005 The Mauthner cell half a century later: a neurobiological model for decision-making? Neuron 47: 13–28.

Lee, Y., Y. Zhang, H. Kang, G. Bang, Y. Kim et al., 2020 Epilepsy- and intellectual disability-associated CYFIP2 interacts with both actin regulators and RNA-binding proteins in the neonatal mouse forebrain. Biochem Biophys Res Commun 529: 1–6.

Lee, Y., Y. Zhang, J. R. Ryu, H. R. Kang, D. Kim et al., 2019 Reduced CYFIP2 Stability by Arg87 Variants Causing Human Neurological Disorders. Ann Neurol 86: 803–805.

Marsden, K. C., R. A. Jain, M. A. Wolman, F. A. Echeverry, J. C. Nelson et al., 2018 A Cyfip2-Dependent Excitatory Interneuron Pathway Establishes the Innate Startle Threshold. Cell Rep 23: 878–887.

Nakashima, M., M. Kato, K. Aoto, M. Shiina, H. Belal et al., 2018 De novo hotspot variants in CYFIP2 cause early-onset epileptic encephalopathy. Ann Neurol 83: 794–806.

Nieuwkoop, P. D., and J. Faber, 1994 Normal Table of Xenopus laevis (Daudin): a systematical and chronological survey of the development from the fertilized egg till the end of metamorphosis.

OMIM, Online Mendelian Inheritance in Man, OMIM®, pp., edited by J. H. U. B. McKusick-Nathans Institute of Genetic Medicine, MD), .

Ortiz, Y. T., L. R. McMahon and J. L. Wilkerson, 2022 Medicinal Cannabis and Central Nervous System Disorders. Front Pharmacol 13: 881810.

Oz, M., K. S. Yang and M. O. Mahgoub, 2022 Effects of cannabinoids on ligand-gated ion channels. Front Physiol 13: 1041833.

Pagano, C., G. Navarra, L. Coppola, G. Avilia, M. Bifulco et al., 2022 Cannabinoids: Therapeutic Use in Clinical Practice. Int J Mol Sci 23.

Panthi, S., P. Chapman, P. Szyszka and C. W. Beck, 2022 Characterisation and automated quantification of induced seizure-related behaviours in *Xenopus laevis* tadpoles. bioRxiv: 2022.2011.2018.517140.

Peng, J., Y. Wang, F. He, C. Chen, L. W. Wu et al., 2018 Novel West syndrome candidate genes in a Chinese cohort. CNS Neurosci Ther 24: 1196–1206.

Sadleir, L. G., E. I. Mountier, D. Gill, S. Davis, C. Joshi et al., 2017 Not all SCN1A epileptic encephalopathies are Dravet syndrome: Early profound Thr226Met phenotype. Neurology 89: 1035–1042.

Scheffer, I. E., S. Berkovic, G. Capovilla, M. B. Connolly, J. French et al., 2017 ILAE classification of the epilepsies: Position paper of the ILAE Commission for Classification and Terminology. Epilepsia 58: 512–521.

Scheffer, I. E., and J. Liao, 2020 Deciphering the concepts behind “Epileptic encephalopathy” and “Developmental and epileptic encephalopathy”. Eur J Paediatr Neurol 24: 11–14.

Schenck, A., B. Bardoni, C. Langmann, N. Harden, J. L. Mandel et al., 2003 CYFIP/Sra-1 controls neuronal connectivity in Drosophila and links the Rac1 GTPase pathway to the fragile X protein. Neuron 38: 887–898.

Schoonheim, P. J., A. B. Arrenberg, F. Del Bene and H. Baier, 2010 Optogenetic localization and genetic perturbation of saccade-generating neurons in zebrafish. J Neurosci 30: 7111–7120.

Sega, A. G., E. K. Mis, K. Lindstrom, S. Mercimek-Andrews, W. Ji et al., 2019 De novo pathogenic variants in neuronal differentiation factor 2 (NEUROD2) cause a form of early infantile epileptic encephalopathy. J Med Genet 56: 113–122.

Session, A. M., Y. Uno, T. Kwon, J. A. Chapman, A. Toyoda et al., 2016 Genome evolution in the allotetraploid frog Xenopus laevis. Nature 538: 336–343.

Sillar, K. T., and R. M. Robertson, 2009 Thermal activation of escape swimming in post-hatching Xenopus laevis frog larvae. J Exp Biol 212: 2356–2364.

Suetsugu, S., D. Yamazaki, S. Kurisu and T. Takenawa, 2003 Differential roles of WAVE1 and WAVE2 in dorsal and peripheral ruffle formation for fibroblast cell migration. Dev Cell 5: 595–609.

Suls, A., J. A. Jaehn, A. Kecskes, Y. Weber, S. Weckhuysen et al., 2013 De novo loss-of-function mutations in CHD2 cause a fever-sensitive myoclonic epileptic encephalopathy sharing features with Dravet syndrome. Am J Hum Genet 93: 967–975.

Ta, A. C., L. C. Huang, C. R. McKeown, J. E. Bestman, K. Van Keuren-Jensen et al., 2022 Temporal and spatial transcriptomic dynamics across brain development in Xenopus laevis tadpoles. G3 (Bethesda) 12.

Trowe, T., S. Klostermann, H. Baier, M. Granato, A. D. Crawford et al., 1996 Mutations disrupting the ordering and topographic mapping of axons in the retinotectal projection of the zebrafish, Danio rerio. Development 123: 439–450.

Vadlamudi, L., L. M. Dibbens, K. M. Lawrence, X. Iona, J. M. McMahon et al., 2010 Timing of de novo mutagenesis--a twin study of sodium-channel mutations. N Engl J Med 363: 1335–1340.

Weuring, W. J., I. Dilevska, J. Hoekman, J. van de Vondervoort, M. Koetsier et al., 2021 CRISPRa-Mediated Upregulation of scn1laa During Early Development Causes Epileptiform Activity and dCas9-Associated Toxicity. CRISPR J 4: 575–582.

Yoo, Y., J. Jung, Y. N. Lee, Y. Lee, H. Cho et al., 2017 GABBR2 Mutations Determine Phenotype in Rett Syndrome and Epileptic Encephalopathy. Annals of Neurology 82: 466–478.

Zhang, Y., H. Kang, Y. Lee, Y. Kim, B. Lee et al., 2018 Smaller Body Size, Early Postnatal Lethality, and Cortical Extracellular Matrix-Related Gene Expression Changes of Cyfip2-Null Embryonic Mice. Front Mol Neurosci 11: 482.

Zhong, M., S. Liao, T. Li, P. Wu, Y. Wang et al., 2019 Early diagnosis improving the outcome of an infant with epileptic encephalopathy with cytoplasmic FMRP interacting protein 2 mutation: Case report and literature review. Medicine (Baltimore) 98: e17749.

Zweier, M., A. Begemann, K. McWalter, M. T. Cho, L. Abela et al., 2019 Spatially clustering de novo variants in CYFIP2, encoding the cytoplasmic FMRP interacting protein 2, cause intellectual disability and seizures. Eur J Hum Genet 27: 747–759.

