## supplemental figures for "Expression of mRNA encoding two gain-of-function *cyfip2* variants associated with DEE65 results in spontaneous seizures in *Xenopus laevis* tadpoles"

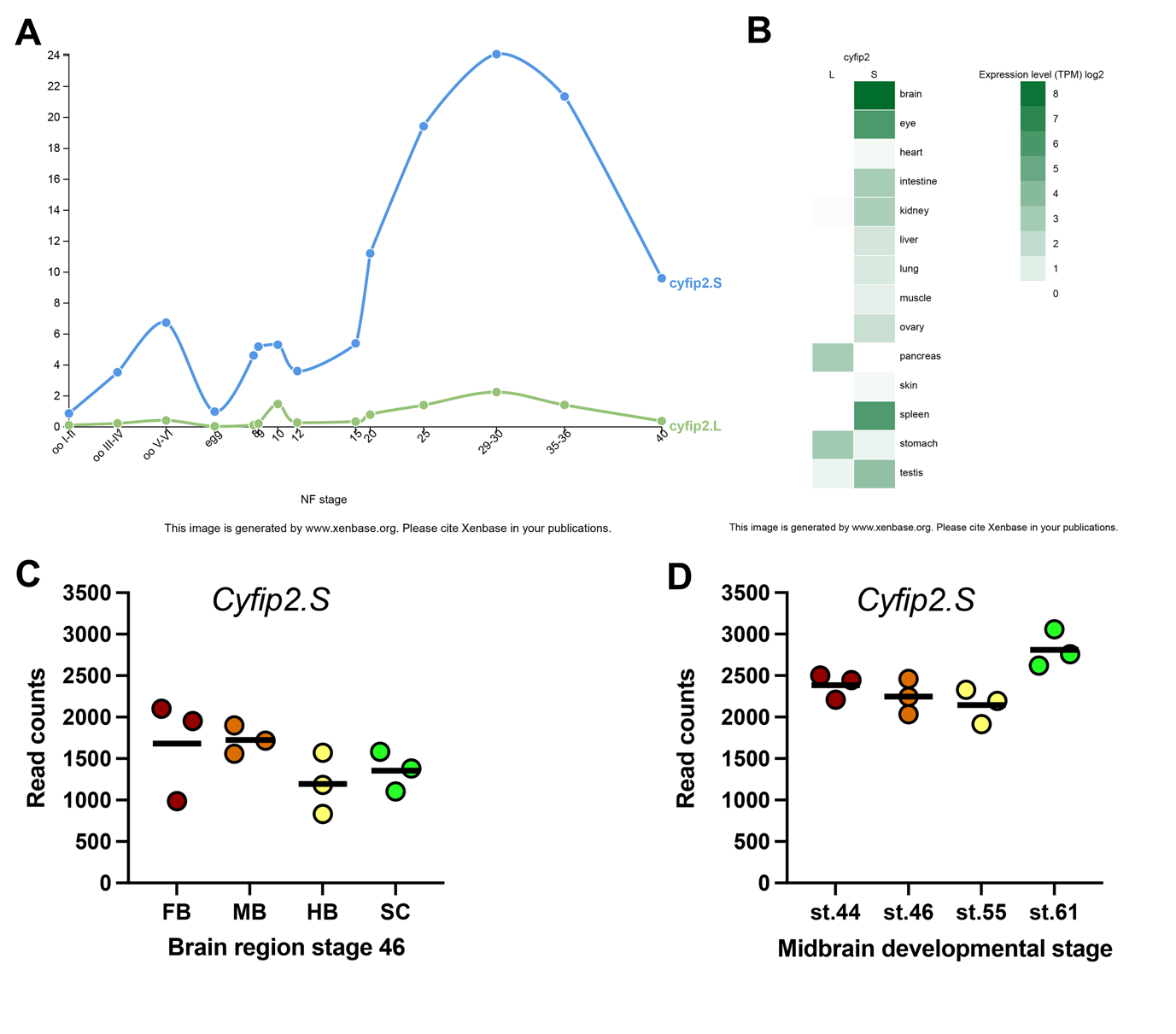

**Figure S1: *Cyfip2.S* is the major homeologue in *X. laevis* and is highly expressed in brain** A,B) Data from Xenbase.org, showing the relative expression of *cyfip2.S* and *L* homeologues across developmental time (A) and across organs (B). C,D) RNAseq Data from Ta et al, 2022, retrived from NCBI GEO, shows high levels of c*yfip2.S* in tadpole brains across three replicate datasets.

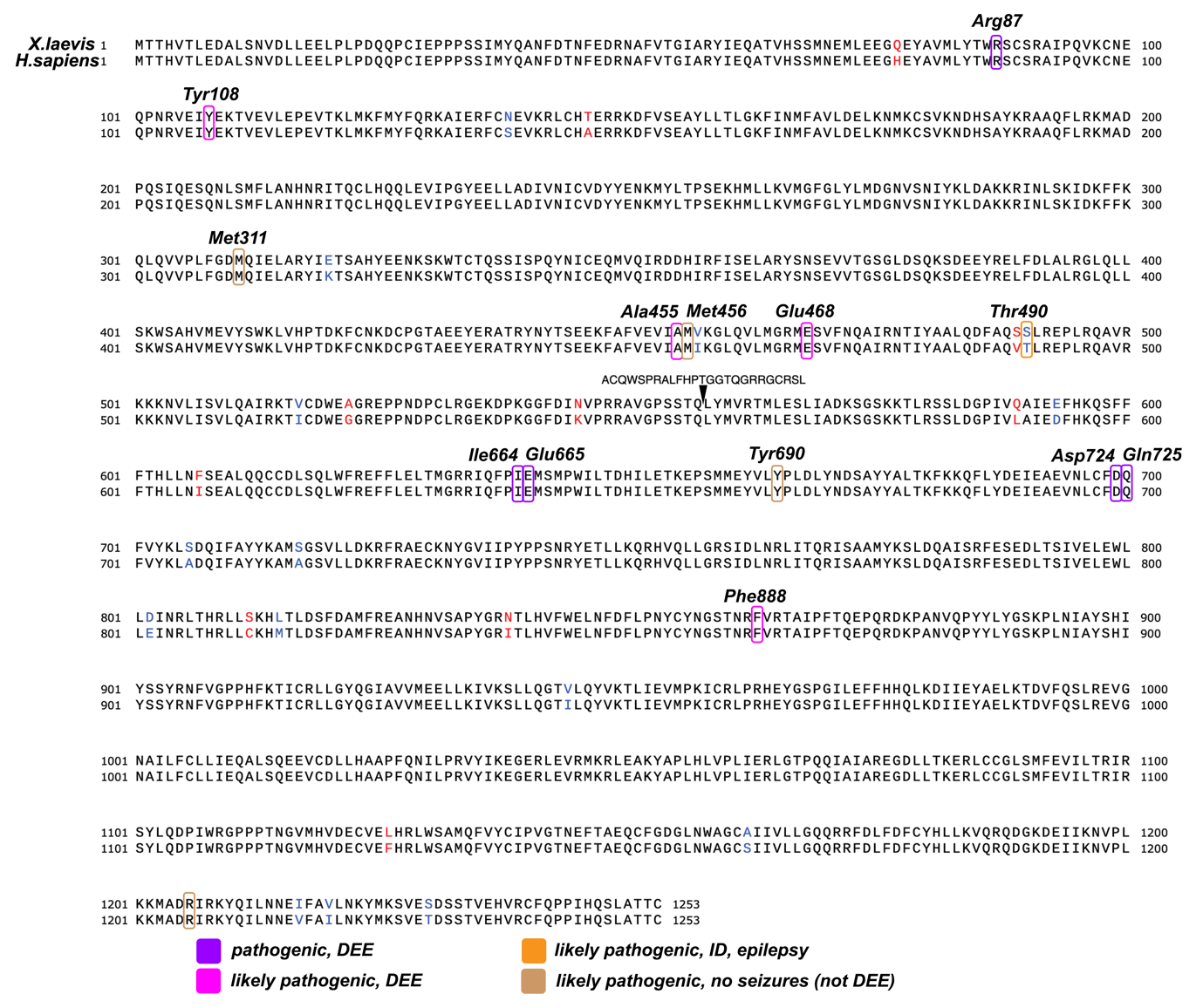

**Figure S2: Protein alignment of *X.laevis* Cyfip2 (top) to *H.sapiens* CYFIP2**, using Smith-Waterman algorithm. Identity is 98%, similarity 99.2%. Arrow indicates location of an exon not annotated in the *Xenopus* sequence. Identical bases are in black, similar bases in blue, and not similar in red. Identified pathogenic and likley pathogenic missense variant positions are shown by coloured boxes, classified as in Begemann, 2021.

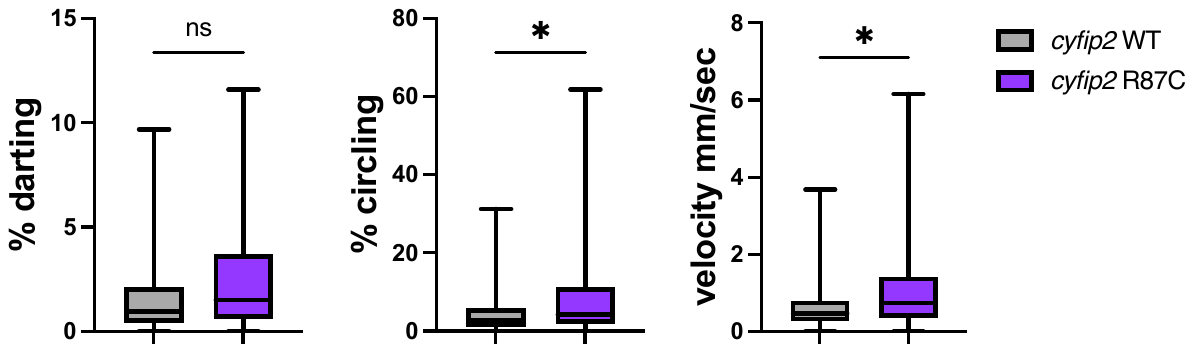

**Figure S3: Replication of *cyfip2*-R87C behavioural phenotype, in an independant cohort of tadpoles at stage 47.** Data from 90 minutes. Box plots, middle line indicates median, box indicates interquartile range and whiskers indicate min/max values, N=24 each group. Unpaired T-test, ns= non-significant, * p<0.05.

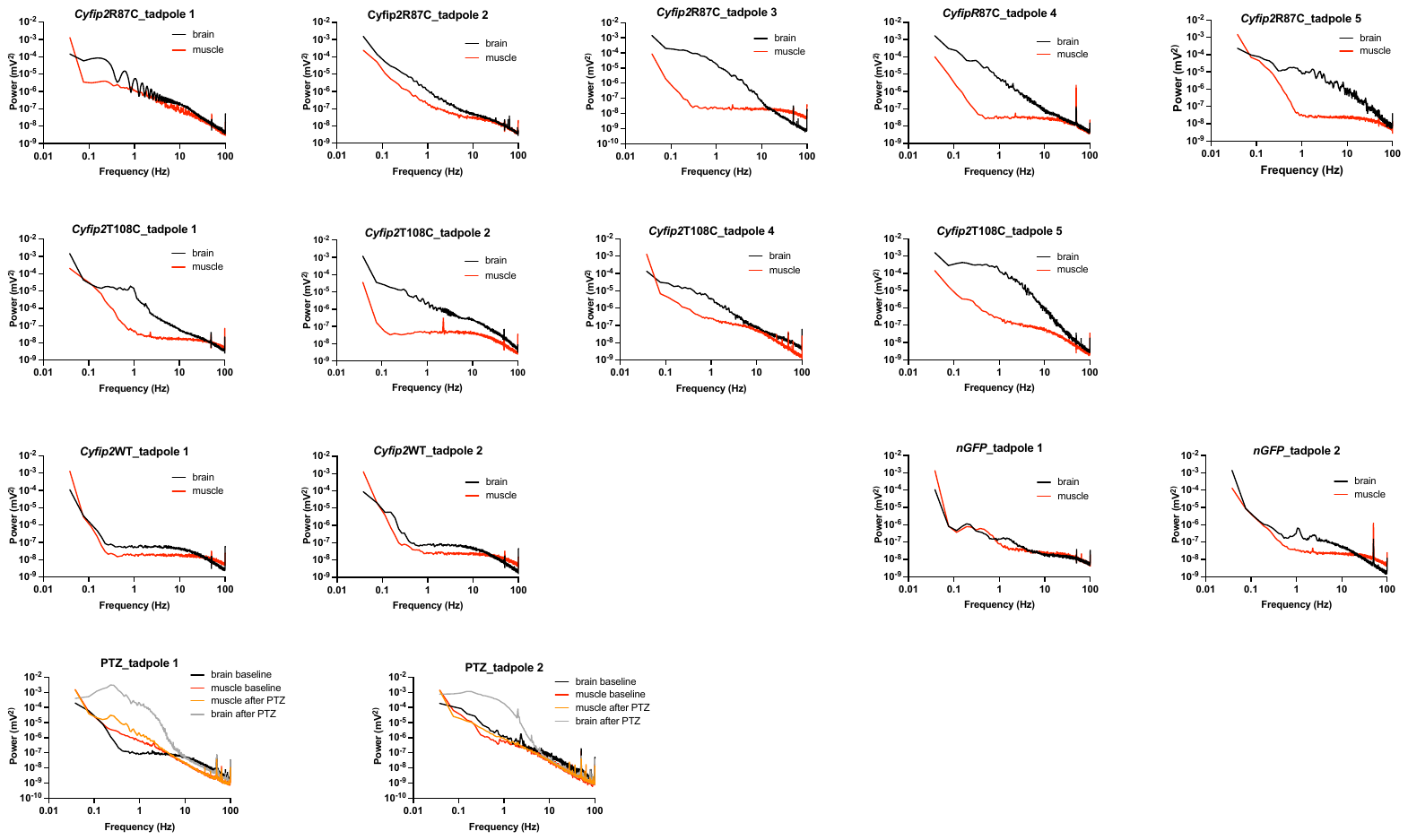

**Figure S4: Power spectra analysis for all tadpoles reported in this study.** This data supports main text figures 3 and 4, generated from Spike 2 v8 and graphed using Prism v9.0.
